# Selective Germline Genome Edited Pig Meninges Grafts for the Abdominal Wall Closure in Damage Control Surgery

**DOI:** 10.1101/2020.02.12.946178

**Authors:** Lijin Zou, Youlai Zhang, Ying He, Hui Yu, Jun Chen, Delong Liu, Sixiong Lin, Manman Gao, Gang Zhong, Weicheng Lei, Guangqian Zhou, Xuenong Zou, Kai Li, Yin Yu, Gaofeng Zha, Linxian Li, Yuanlin Zeng, Jianfei Wang, Gang Wang

## Abstract

Reconstruction of abdominal wall defects is still a big challenge in surgery, especially where there is insufficient fascia muscular or excessive tension of the defects in emergency and life-threatening scenarios. Indeed, the concept of damage control surgery has been advanced in the management of both traumatic and nontraumatic surgical settings. The strategy requires abridged surgery and quick back to intensive care units (ICU) for aggressive resuscitation. In the damage control laparotomy, patients are left with open abdomen or provisional closure of the abdomen with a planned return to the operating room for definitive surgery. So far, various techniques have been utilized to achieve early temporary abdominal closure, but there is no clear consensus on the ideal method or material for abdominal wall reconstruction. We recently successfully created the selective germline genome-edited pig (SGGEP) and here we aimed to explore the feasibility of in vivo reconstruction of the abdominal wall in a rabbit model with SGGEP meninges grafts (SGGEP-MGs). Our result showed that the SGGEP-MGs could restore the integrity of the defect very well. After 7 weeks of engraftment, there was no sign of herniation observed, the grafts were re-vascularized, and the defects were well repaired. Histologically, the boundary between the graft and the host was very well integrated and there was no strong inflammatory response. Therefore, this kind of closure could help restore the fluid and electrolyte balance and to dampen systemic inflammatory response in damge control surgery while ADM graft failed to establish re-vascularization as the same as the SGGEP-MG. It is concluded that the meninges of SGGEP could serve as a high-quality alternative for restoring the integrity of the abdominal wall, especially for damage control surgery.

## Introduction

The congenital and acquired abdominal wall defects due to trauma, operation, tumour resection pose a formidable hurdle to the surgeons. Many different methods have been used for reconstruction but the optimal method is yet to be found. With the advent of prosthetic meshes the hernia recurrence rate is reduced (1, 2), but these implants could not establish re-vascularization and are not bioabsorbable. They not only could potentially cause the infection, chronic pain but also lead to other organ dysfunction, for example, intestine adherence or obstruction (3–10).

Notably, in the emergency and life-threatening scenario, the immediate goal is the closure of fascial defect as early as possible without triggering or exacerbating the lethal abdominal compartment syndrome (11–13). However, the management of these patients gives rise to a daunting challenge for surgeons. So far, techniques such as packing, mesh, and vacuum-assisted closure have been utilized to achieve early temporary abdominal closure. Nevertheless, there is no clear consensus on the ideal method or material for abdominal wall reconstruction. Each technique or material has their advantage and disadvantage(14, 15).

Very recently, we successfully established the selective germline genome edited pig (SGGEP) with a better immunological and molecular compatibility profile for humans (https://doi.org/10.1101/2020.01.20.912105). In this pig, three xenoantigens (Glycoprotein alpha-galactosyltransferase 1, GGTA;β-1,4-N-acetyl-galactosaminyl transferase 2, B4GAL and, cytidine monophosphate- N- acetylneuraminic acid hydroxylase, CAMH) which are well known to cause hyperacute rejection in xenotransplantation are knocked out, and additional eight human immune compatibility enhanced genes are knocked in, these genes encode a cluster human proteins including human complement system negative regulatory proteins (CD46, CD55, and CD59), human coagulation system negative regulatory proteins thrombomodulin (THBD), tissue factor pathway inhibitor (TFPI); CD39; macrophage negative regulatory proteins (human CD47); and natural killer cell negative regulatory human leukocyte antigen class I histocompatibility antigen, alpha chain E (HLA-E). When SGGEP skin was engrafted to NHP, up to 25 days’ graft survival was observed, thus SGGEP skin grafts demonstrated the long immunological tolerance in NHP (https://doi.org/10.1101/2020.01.20.912105).

In the present study, we aimed to extend the utility for our SGGEP, we set out to explore the feasibility of in vivo reconstruction of the abdominal wall in a rabbit model using SGGEP meninges grafts (SGGEP-MG). Our results indicated that the SGGEP-MGs could repair the abdominal wall defect well by establishing revascularization and restoring the integrity of defected abdominal wall after 7 weeks engraftment while the pig acelluar Derm Matrix(ADM) grafts failed to repair the defects. Therefore, SGGEP-MG can be considered as a high-quality alternative, specially for the temporary closure of fascial defect techniques in the damage control surgery. This encouraging data offers the justification and a conceptual framework for ongoing NHP preclinical safety and efficacious assessment and potential future clinical applications.

## Results

The clinical gross appearance of the grafts was assessed every 7 days after transplantation. The post-operation day 49 was designed as the endpoint in the experiment group while in the control group, the post-operation day 35 was set as the endpoint. It was observed that the SGGEP-MG repaired abdominal wall defects well, it offered good healing for the defect with collagen fibrils-rich environment (Figure2 d, e, f). There was no sign of herniation throughout the experiments observed. Importantly, the grafts were-vascularized and integrated with surrounding tissues very well, there was no strong inflammation response (Figure1e,f,g,h) throughout the whole course of the study. The histological analysis confirmed these observations with minor inflammatory cell cells and rich of collagen in both junctional and central regions in the grafts (Figure2 d, e, f). The collagen fibres were still observed by 7 weeks after transplantation and high-density vessels were evidenced (Figure 2 d, e, f).

**Figure 1.**
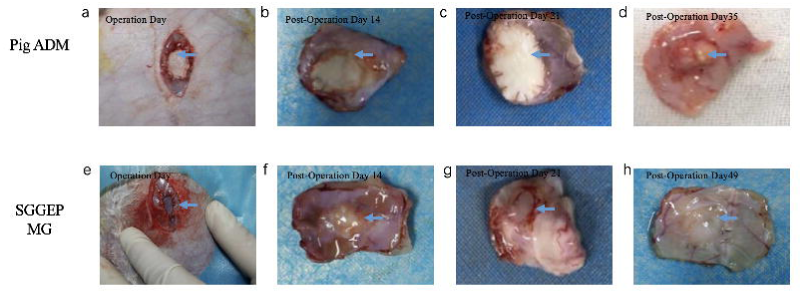
The respective gross appearance picture were shown after Selective Germline Genome Edited Pig meninges grafts(SGGEP-MG) to rabbit peritoneum xenotransplantation. (a) Pig ADM(acellular dermal matrix) graft on the operation Day, (b) Pig ADM graft on the post-operation Day14,(c) Pig ADM graft on the post-operation Day 21,(d) Pig ADM graft on the post-operation Day 35,(e) SGGEP meninges(SGGEP-MG) graft on the operation Day(f) SGGEP-MG on the post-operation Day 14,(g) SGGEP-MG on the post-operation Day 21,(h) SGGEP-MG on the post-operation Day49.

**Figure 2.**
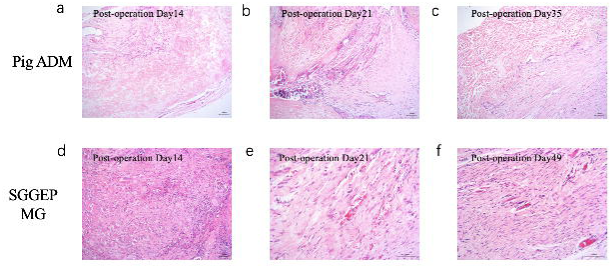
The respective histology HE staining images are shown from biopsies taken on after Selective Germline Genome Edited Pig meninges grafts(SGGEP-MG) to rabbit peritoneum xenotransplantation. (a) Pig ADM graft on the post-operation Day 14,(b) Pig ADM graft on the post-operation Day21,(c) Pig ADM graft on the post-operation Day 35,(d) SGGEP-MG on the post-operation Day 14,(e) SGGEP-MG on the post-operation Day 21,(f) SGGEP-MG on the post-operation Day 49

In the control group, ADM grafts yielded quite different graft-host healing responses although there was also no sign of herniation observed. The boundary between the graft-host was clear both at clinical observation and histological analysis (Figure 1 a, b, c, d, Figure 2 a, b, c) indicating that the grafts were not re-vascularized and integrated with neighbouring host tissues. The HE staining indicated that though the ADM could provide the temporary cover for the defect but failed to repair the abdominal wall defect by establishing revascularization and restoring the integrity as quickly as the SGGEP group.

## Materials & Methods

### Animals

All studies were approved by the First Affiliated Hospital of Nanchang University Institutional Animal Care and Use Committee (IACUC) and performed in accordance with the Guidelines for the Humane Treatment of Laboratory Animals (2006) issued by the Chinese Government.

(http://www.most.gov.cn/fggw/zfwj/zfwj2006/200609/t20060930_54389.htm. Japanese white rabbits (2.5 ± 0.5kg, 6-month-old, male; Longping Company, Nanchang, China) and SGGEP (~3 months old) were used in these studies.

### SGGEP meninges grafts xenotransplantation in a rabbit peritoneum defect model

Swine donors were anesthetized with 2.5mg/kg diazepam IM and 10mg/kg ketamine IM. Then, endotracheal intubation was performed for anesthesia (2% isoflurane and oxygen). The head was shaved using clippers and then disinfected with Povidone-Iodine and rubbed with 70% ethanol, the skull was opened, 5 × 5 cm cerebral dura grafts were harvested, the grafts for immediate use were immersed in saline and stored at 4°C. The pigs were all euthanized by an overdose of anesthetics.

The rabbits were anesthetized with 10% chloral hydrate (4 ml/kg) through intraperitoneal injections. The animals were divided into two groups. The experimental group (n = 7) were received SGGEP cerebral dura grafts. The control group (n=5) were received ADM (acellular dermal matrix) grafts. A longitudinal skin incision was made and abdominal wall muscle was dissected bluntly. A 5×5 cm peritoneal defect was made carefully by the ophthalmic scissor. For the experimental group, SGGEP-MGs were implanted to the defect site appropriately to close the defect while ADM was implanted as a control. The grafts were sutured to rabbit abdominal defect with 3–0 Ethilon by interrupted sutures.

Pressure bandaging was applied to protect the incisional site. Ceftriaxone sodium (80mg/kg) IV was administered once before and once after the operation. All rabbits were individually housed in a cage after the surgery. The graft clinical assessment was made as indicated time points to analyze the graft-host healing effects.

### Histopathology analysis

After the engraftments, the samples were collected on different days. The samples were fixed in 4% paraformaldehyde and embedded in paraffin. Sections (5µm) were cut, mounted on the slide, and stained with hematoxylin and eosin (HE). The HE sections were viewed under x 100 magnification, photos were taken for analysis.

## Discussion

To close the abdominal wall, various techniques have been used for the past century for different etiology, location and degree (layers implicated) of the defects. Some utilized the mesh (16–18), the others used the flaps (19–21). Recent new procedures with better results were reported (22–24). However, each technique has its advantage and disadvantage. Ideally, after implantation of the mesh, several key host processes should occur, including the acute inflammatory response, followed by the migration of mononuclear cells to the site of implantation. After cellular penetration of the material, new blood vessel proliferation (neovascularization) extends through the thickness of the matrix. Concurrently, mononuclear cells (macrophages) secrete cytokines and other signalling molecules, resulting in the recruitment of fibroblasts to the region, deposition of endogenous collagen, and resorption of the mesh. These processes are considered integral to biologic mesh integration and remodelling (25).

With the aim of obtaining a tension-free closure of the abdominal wall defects, synthetic materials are often utilized to restore missing fascial tissues and strengthen the healing. But these nonabsorbable materials may cause complications, for instance, infections, chronic pain, erosions, and fistula formation, in particular when they are set directly over the viscera. While other materials are available, but each has its own pros and cons (26–28).

Especially, the damage control surgery practice is one of the greatest progress in recent decades and has established as a common procedure in both traumatic and general surgery. One of the damage control surgery purposes is the closure of the fascial defect as quickly as is clinically possible without rising the intraabdominal pressure during the initial hospitalization. Quite a few techniques have been reported workable in helping to increase the primary abdominal fascial closure rate. However, currently, high-quality comparative data is rare (15).

Biological materials have an advantage over these nonabsorbable and no blood vessel materials in view of maintaining physiological homeostasis. Very recently we establish the SGGEP with enhanced human compatibility (https://doi.org/10.1101/2020.01.20.912105). We aimed to evaluate if SGGEP MGs, rich in collagen and strong tensile strength, could serve as a better alternative. Moreover, we would like to assess if it could fulfill the re-vascularization which is a key to maintain the water and electrolyte balance and to combat the infection in the damge control surgery.

Indeed, in the present study, after implantation into abdominal wall defect, SGGEP MGs were observed to be re-vascularized, maintain its structural integrity and the boundary between the graft and the surrounding tissues was integrated indicating a good host –graft environment. While the ADM graft did not yield a good host-graft interaction and the boundary between the graft and host was clearly demarcated throughout the experiment course. Considering the effective repair of abdominal wall defects relying on the early re-establishment of vascular vessels (29), the SGGEP graft had a very satisfying result.

As it is well known that an ideal graft should have characteristics such as strong, inert, resistant to infection, low recurrence rate, high quality of life and good value. Ongoing advancements in techniques and technology will continue to help surgeons tailor the graft to the clinical scenario(30). Therefore, it can be concluded that the SGGEP meninges graft is more effective for managing abdominal wall defects than ADM. It has the potential to be widely applied to deal with abdominal wall defects, particularly in emergency or infection situations. This study points out that the SGGEP meninges graft is effective and safe for the reconstruction of abdominal wall defects. SGGEP meninges implantation is a feasible and reproducible technique providing a large and functional covering of abdominal wall defect. The SGGEP meninges can be cryopreserved, stored for emergency events, and transported worldwide for global use (Figure 3). Further NHP pre-clinical investigations are necessary to assess the practical validity of this concept.

**Figure 3.**
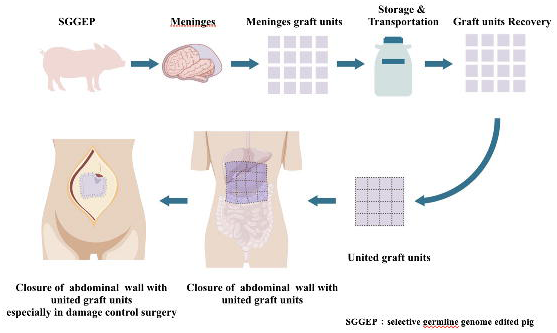
Working model for the SGGEP meninges grafts applications

## Acknowledgements

L.Z., Y.Z. was supported by the grant Natural Science Foundation of China(NSFC 81660364 and NSFC 81760343).X.Z. was funded by grants Natural Science Foundation of Guangdong Province (2017A030308004) and Natural Science Foundation of Guangzhou City (201804020011). G.Z. was supported by the National Key R&D Program of China (2017FA105202) and Shenzhen Scientific & Innovation Committee (GJHZ20180928155604671),Y.H. was funded by Health and Family Planning Project of Jiangxi Province(No. 20191025).

## Competing Financial Interests

J.W., G.W., K.L. are inventors on a patent filed by Geneo Medicine Inc using the technology described in this paper. J.W, G.W, K.L., are employees of Geneo Medicine Inc, G.W.,W.L is an employee of Zhuzao Biotechology, Inc.

